# Surface morphometrics reveals local membrane thickness variation in organellar subcompartments

**DOI:** 10.1101/2025.04.30.651574

**Authors:** Michaela Medina, Ya-Ting Chang, Hamidreza Rahmani, Daniel Fuentes, Benjamin A. Barad, Danielle A. Grotjahn

## Abstract

Lipid bilayers form the basis of organellar architecture, structure, and compartmentalization in the cell. Decades of biophysical, biochemical, and imaging studies on purified or *in vitro* reconstituted liposomes have shown that variations in lipid composition influence the physical properties of membranes, such as thickness and curvature. However, similar studies characterizing these membrane properties within the native cellular context have remained technically challenging. Recent advancements in cellular cryo-electron tomography (cryo-ET) imaging enable high-resolution, three-dimensional views of native organellar membrane architecture preserved in near-native conditions. We previously developed a ‘Surface Morphometrics’ pipeline that generates surface mesh reconstructions to model and quantify cellular membrane ultrastructure from cryo-ET data. Here, we expand this pipeline to measure the distance between the phospholipid head groups (PHG) of the membrane bilayer as a readout of membrane thickness. Using this approach, we demonstrate thickness variations both within and between distinct organellar membranes. We also demonstrate that membrane thickness positively correlates with other features, such as membrane curvedness. Further, we show that subcompartments of the mitochondrial inner membrane exhibit varying membrane thicknesses that are independent of network morphology (i.e., fragmented versus elongated networks). Finally, we demonstrate that large membrane-associated macromolecular complexes exhibit distinct density profiles that correlate with local variations in membrane thickness. Overall, our updated Surface Morphometrics pipeline provides a framework for investigating how changes in membrane composition in various cellular and disease contexts affect organelle ultrastructure and function.

## Introduction

Eukaryotic cells rapidly remodel their organellar membranes in response to a variety of cellular stresses and physiological conditions. Lipid bilayers, the core unit of organellar membranes, are created through the self-assembly of amphipathic phospholipids that orient their hydrophilic head groups to shield their hydrophobic tails from the aqueous cytoplasmic environment. Their dynamic and fluid nature enables membrane remodeling required for many diverse cellular processes, including inter-organellar communication, scaffolding, organelle biogenesis, and quality control. In addition to phospholipids, integral membrane proteins also dictate the inherent properties of organellar membranes. Membrane proteins such as ATP synthase dimers have the capacity to deform, bend, and stabilize the phospholipid bilayer^1^. Conversely, membrane properties such as stiffness and curvature can impact the folding, localization, and function of integral membrane proteins^2^. Through dynamic remodeling of lipid and membrane protein composition, organelles can modulate their shape and thus function to facilitate cellular processes, such as vesicular trafficking^3,4^, inter-organellar resource transfer^5^, and organellar fission and fusion^6,7^. Despite this important link between membrane remodeling and adaptive function, defining local changes to lipids and proteins in their native context has remained technically challenging.

Under conditions of low dose and defocus^8^, cryo-electron microscopy (cryo-EM) imaging has the resolving power to distinguish the two opposing rows of phospholipid head groups that comprise the lipid bilayer and the proteins embedded within them. Several groups have harnessed the power of cryo-EM to reveal how different compositions of phospholipids, when assembled in giant unilamellar vesicles, can impart distinct membrane properties, including membrane thickness, rigidity, and compressibility^8–10^. In combination with cell thinning techniques such as cryo-focused ion beam milling, it is now possible to obtain high-resolution views of lipid bilayers within their native cellular context using cryo-electron tomography (cryo-ET) imaging. There are also multiple voxel-based density sampling strategies developed to analyze protein structure and reveal the organization of the embedded proteins within membranes visible in cellular cryo-ET data^11,12^. However, to date, none of these density-sampling approaches have been adapted to calculate membrane thickness directly and specifically. A recently reported voxel segmentation based approach^13^ estimates membrane thickness by generating and converting voxel segmentations of membranes into oriented point clouds to approximate the edge points as the membrane “boundaries”. Ray projections between opposing boundary edge points are then used as a proxy to estimate membrane thickness. While this approach revealed relative changes in thickness, the approach led to measurements considerably larger than previously reported for *in vitro* vesicles, and an approach that allows correlation with other features of membrane geometry would be beneficial.

Here we present a new method for measuring membrane thickness using triangulated surface mesh reconstructions to calculate voxel density-based line scans across organellar membranes. Surface mesh reconstructions provide a more accurate model of the inherent geometry of the membrane, independent of voxel size. By calculating per-triangle density line scans across these surface mesh models, we generate plots of local membrane density extracted from the tomogram itself. We demonstrate that the opposing phospholipid head groups of the membrane bilayer result in peaks within these line scans; we use the distance between these peaks as the membrane thickness, consistent with previous studies^9^. This model-guided approach also integrates with other ultrastructural measurements in the Surface Morphometrics pipeline, enabling correlation with geometric features such as curvature and inter-membrane spacing. We show that this approach can be used to detect statistically significant differences in membrane thickness across organelles and their functionally distinct membrane compartments. Furthermore, we demonstrate that large membrane-associated macromolecular complexes exhibit distinct density profiles that correlate with local variations in membrane thickness. By integrating these density-based measurements within our existing Surface Morphometrics pipeline^14^, we provide the first automated approach to measure intracellular membrane thickness and correlate it with the rest of the cellular context.

## Results

### Measuring organelle membrane thickness in cellular cryo-electron tomograms

We set out to develop an accurate and automated pipeline for measuring organellar membrane thickness from cellular cryo-ET data directly (***Figure 1***). In brief, we performed cryo-focused ion beam milling to generate thin (∼80-150 nm) cellular sections (i.e., lamella) of vitrified mouse embryonic fibroblasts with mitochondria-targeted GFP (MEFmtGFP) cultured on electron microscopy grids. To resolve the leaflet bilayer of organelles, we used low-to-medium defocus (4-6 μm) and low dose cryo-ET imaging to acquire tilt series of regions primarily containing mitochondria and other associated organelles. This defocus range maximizes contrast while minimizing the defocus-based blurring between bilayers that leads to poor fits. We reconstructed the resulting tilt series into tomograms and used automated U-net-based methods^15^ to generate binarized membrane segmentations, which were further labeled to distinguish between cellular membranes such as the outer and inner mitochondrial membranes (OMM and IMM, respectively), endoplasmic reticulum (ER) membranes, and vesicles. The resulting voxel segmentation models were processed through the Surface Morphometrics pipeline^14^ (https://github.com/GrotjahnLab/surface_morphometrics) to generate triangulated surface meshes that approximate the mid-surface between the two sides of the bilayer (***Figure 1A***).

**Figure 1:**
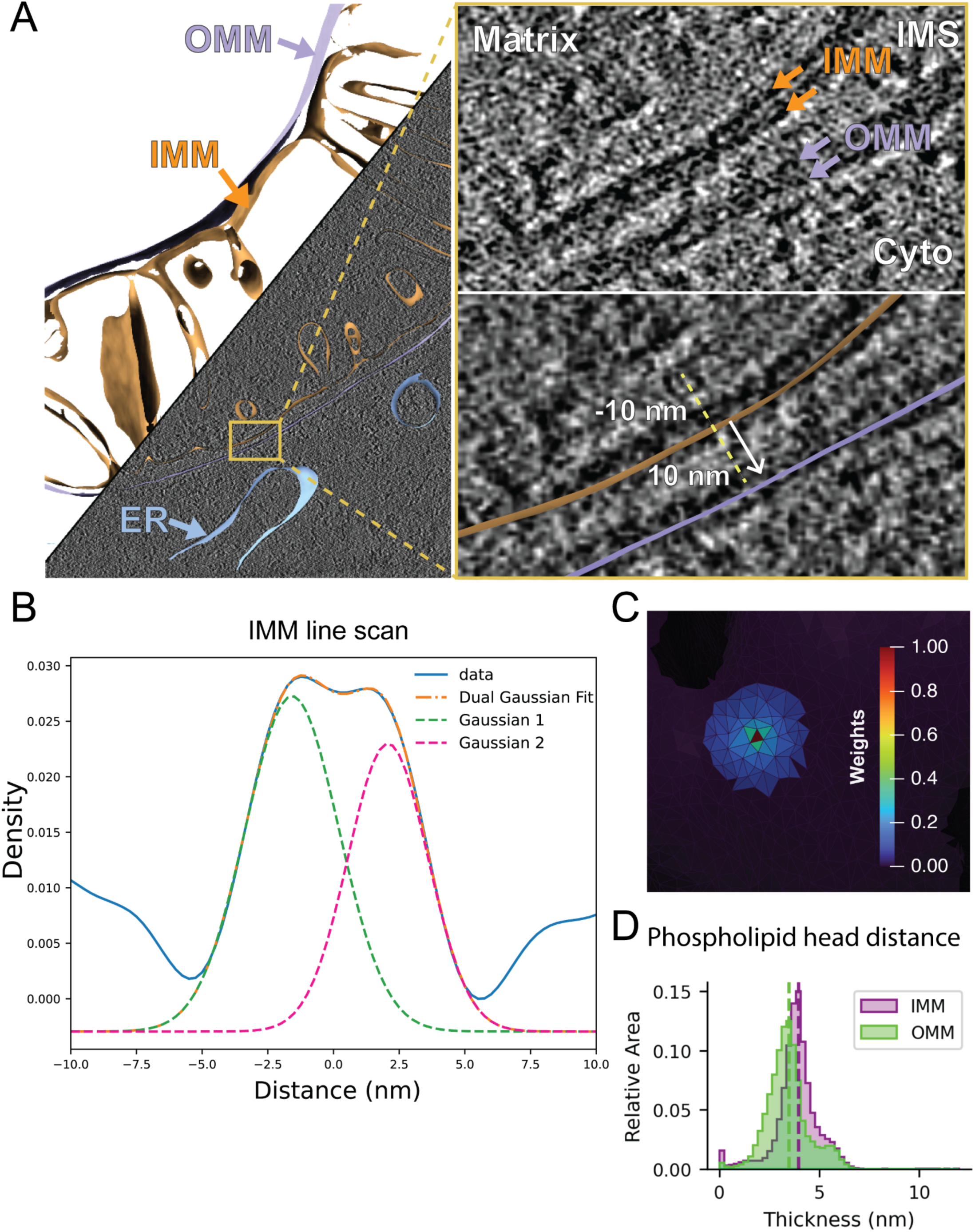
Extending Surface Morphometrics to measure lipid bilayer thickness within tomograms. A. Triangle mesh models for both IMM (orange) and OMM (lavender) are overlaid on top of the binned tomogram reconstruction which is of sufficient resolution to distinguish lipid head groups visually. The meshes can be used to generate local vectors for density line-scan analysis within the tomogram (shown in inset). B. The line scans generated by sampling along these normal vectors can be combined across a surface to generate a smooth plot with peaks corresponding to the two headgroups of the bilayer (Blue line). We fit the curve with a dual Gaussian model (Orange dashed line, individual gaussians pink and green) and used the peak to peak distance as the bilayer distance. C. Local thickness variation analysis was determined via a local weighted averaging scheme, in which the line scan of the central triangle (red) is combined with neighboring triangles up to 12 nm away with weights that decrease with distance from the center. D. Using per-triangle thickness measurements, we identified the differences in the distribution of thicknesses within different membranes, as depicted in a triangle-area-weighted histogram, whose Y-axis measures the relative area of membrane in each thickness bin.

To measure thickness, we start from the midpoint of each triangle on the surface mesh and then systematically interpolate the tomogram density along a 10 nm path, following the normal vector in both the positive and negative directions (***Figure 1A*)**. We use these measurements to generate a voxel line scan, which reveals two peaks corresponding to the densities associated with the phospholipid head groups (PHG) (***Figure 1B***). To estimate the positions of these PHGs, we fit the line scans with a curve composed of two Gaussian distributions and measure the distance between the two Gaussian peaks as a readout of PHG spacing, which is generally used as the metric for membrane thickness^9^. We found that estimating these peaks from non-denoised Warp back-projected tomograms is challenging and prone to artifacts due to the low signal-to-noise ratio. We therefore performed this analysis on binned tomograms (9.98 A/pix) and performed distance-weighted averaging of the signal of triangles within a 12 nm radius (***Figure 1C***), hyperparameters which we determined heuristically had a good balance of local information with robust measurement of thickness. We used this approach to measure PHG distance locally across every triangle of each surface mesh reconstruction in our dataset and show the different PHG distances between different organelles, revealing an apparent increase in thickness in the IMM compared to the OMM (***Figure 1D***). Upon inspection of the generated surfaces, we observed less reliable thickness measurements at the edges of the surface meshes due to artifacts caused by the missing wedge (***Supplemental Figure 1***). To ensure that we only included membranes that had reliable thickness estimations, we implemented an edge exclusion feature to exclude all triangles within 8 nm of the edge of the original surface. This led to robust and accurate membrane thickness measurements that are free from artifacts inherent to cryo-ET data. All reported membrane thickness measurements were generated with edge filtering applied to the surface mesh reconstructions.

### Cellular membranes display significant differences in membrane thickness

Previous biochemical and structural analyses of purified and reconstituted membranes have demonstrated that membranes with unique lipid and membrane compositions result in changes to the biophysical properties of the lipid bilayer, such as thickness^16^. To assess whether these differences are observed in the native cellular environment, we applied our pipeline to calculate membrane thickness on a per-triangle basis across all triangles within each surface mesh for every organellar membrane in our dataset. Plotting the spatial distribution of these thicknesses directly on the surfaces (***Figure 2A, 2B, Supplemental Figure 2***) and the combined distribution of thicknesses for all surface mesh triangles on a histogram (***Figure 2C***) revealed subtle differences in thickness across cellular organelles. Assessing the median thicknesses as individual observations for each organelle showed significant differences in membrane thickness across cellular compartments (***Figure 2D***). Interestingly, our results show that the OMM is significantly thinner (3.2±0.10 nm) compared to the IMM (3.6±0.07 nm), demonstrating that variations in membrane thickness exist even within the same organelle. The OMM also showed statistically significant reductions in membrane thickness relative to ER (3.7±0.05 nm) and vesicle (3.6±0.08 nm) membranes, which were also significantly different from each other. Taken together, we show that different organelle membranes exhibit significant differences in average membrane thickness, demonstrating the power of our approach to quantify subnanometer-level differences in membrane thickness across membranes visualized in their native environment.

**Figure 2:**
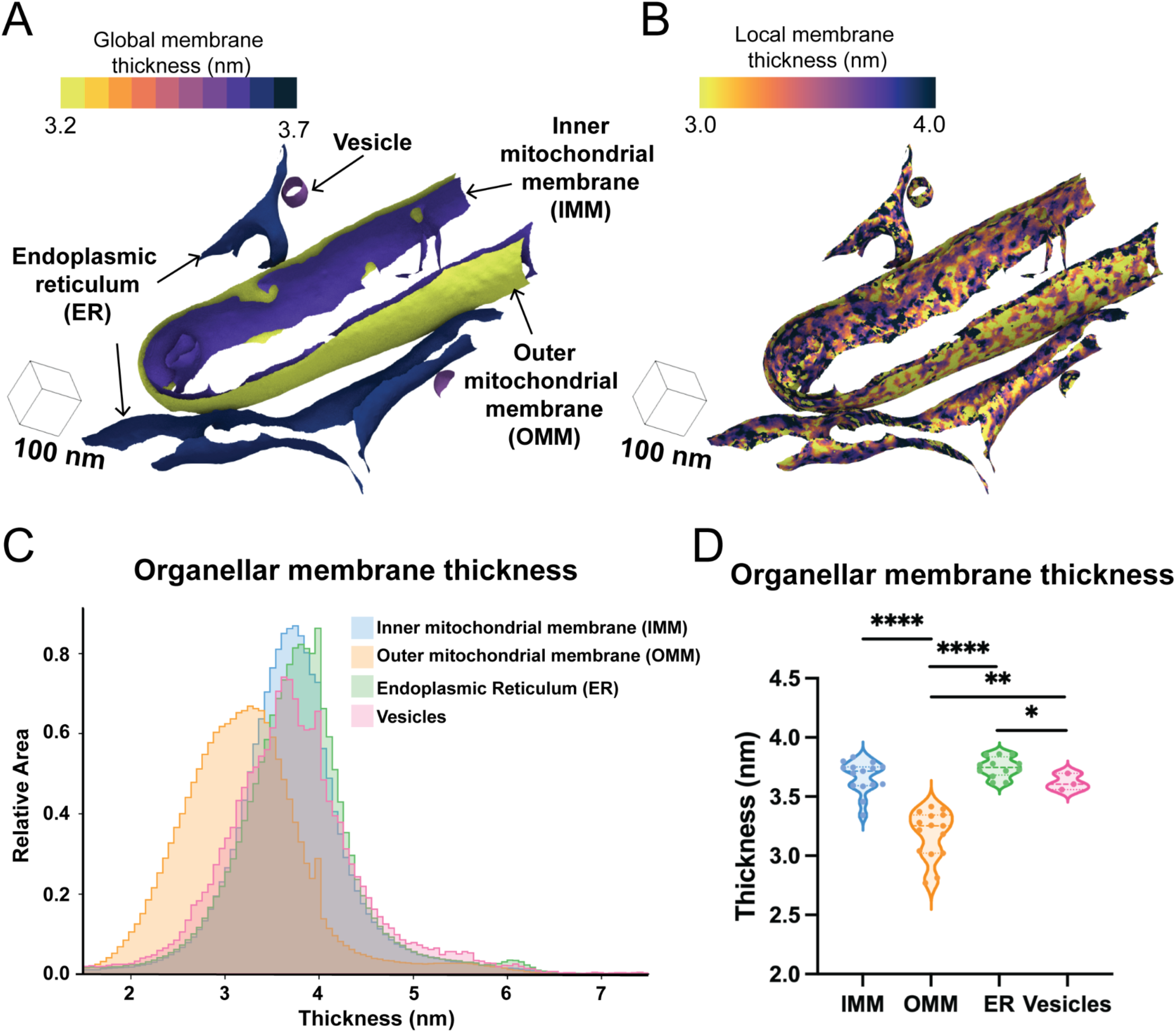
Membrane thickness varies across organelles. A. The global and local per-triangle measurements across different organelle classes are mapped to the different organelle classes observed in a single tomogram. B. The local per-triangle measurements across different organelle classes are mapped to the different organelle classes observed in a single tomogram. C. Per-triangle median thickness histograms highlight the difference in thickness distributions between different physiological organelles. D. Violin plot of per-surface median thickness reveals statistically significant differences in thickness between different organelles. IMM: n = 15; OMM: n =15; ER: n = 12; Vesicle: n =3. P values from Mann-Whitney U test are indicated. *P < 0.05; ** P < 0.01, ****P < 0.001.

### Subcompartments of the inner mitochondrial membrane vary in membrane thickness

We next asked whether variations in membrane thickness are observed across functionally distinct regions within the same organellar membrane. Specialized lipids and proteins influence the shape of the IMM and help fold it into distinct subcompartments, including the regions closely appressed to the OMM, termed the inner boundary membrane, the protruding regions called the cristae body, and the transition zones between these regions, called the cristae junctions. We previously showed that our Surface Morphometrics pipeline can automatically classify between these distinct compartments based on their distance from the OMM^14^. We performed a similar subclassification procedure on the IMM in this dataset and measured the membrane thickness locally within each of these subcompartments (***Figure 3A***). Interestingly, we observed significant differences in the thickness of these compartments, with the cristae body exhibiting significantly thicker membranes (3.8±0.04 nm) relative to the crista junction and inner boundary membrane compartments (***Figure 3B* *& C*)**. While subtle, this significant change in membrane thickness across the contiguous membrane suggests that these differences may reflect an additional structural regulation of IMM subcompartment specialization.

**Figure 3.**
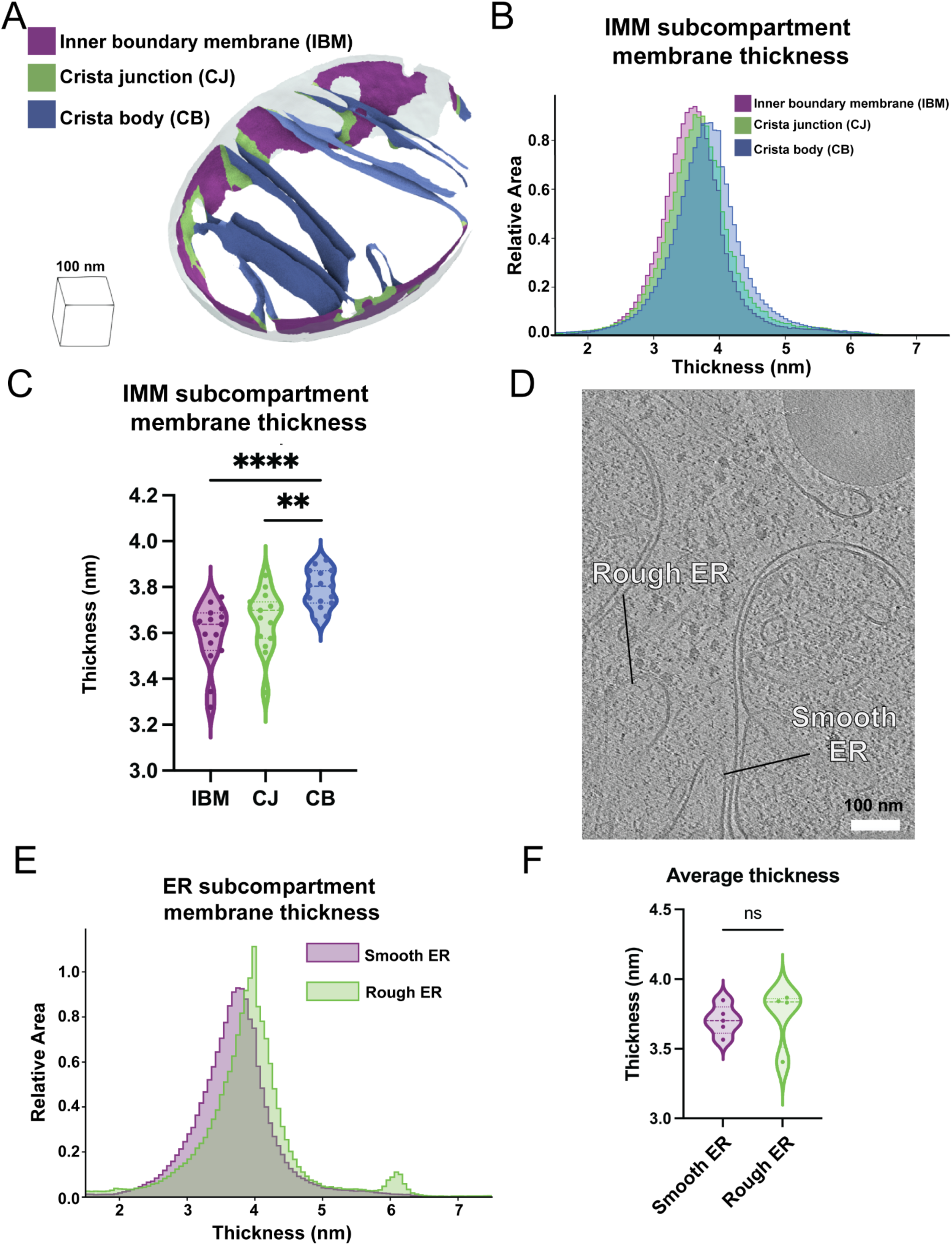
Average membrane thickness varies within subcompartments of the same organelle. A. OMM distance-based classification automatically segments the inner boundary membrane (IBM, purple), crista junctions (CJ, green), and crista bodies (CB, blue) in surface mesh models. B. An area-weighted histogram of per-triangle thickness measurements shows variation between different mitochondrial subcompartments. C. Violin plot of per-tomogram median thickness shows statistically significant variations between mitochondrial subcompartments. IBM: n = 15; CJ: n = 15; CB n = 15. P values from Mann– Whitney U test are indicated. ** P < 0.01, ****P < 0.001. D. Tomogram showing rough and smooth ER. Classification based on the presence or absence of bound ribosomes on the ER membrane, respectively. E. Area-weighted histogram of per-triangle thickness measurement across ER subcompartment. F. Violin plot of per-tomogram median thickness of each ER subcompartment. Rough ER: n = 4; Smooth ER: n = 5.

Like the IMM, the ER membrane can be further subdivided into two spatially and structurally distinct compartments: the rough ER, comprised of sheet-like structures studded with co-translating ribosomes, and the smooth ER, which forms tubular projections that often make functional contacts with other organelles^17,18^. We leveraged these distinct structural characteristics to classify the two types of membranes in our dataset based on the presence of membrane-docked ribosomes (rough ER) or the absence of ribosomes (smooth ER) (***Figure 3D*)**. In contrast to the subcompartments of the IMM, we detected no significant differences between the smooth and rough ER, with membrane thicknesses of 3.7±0.09 nm and 3.7±0.22 nm, respectively. (***Figure 3E* *& F***).

### Membrane thickness positively correlates with membrane curvature

Given the observed IMM compartment-specific differences, we wondered whether other aspects of mitochondrial structure are associated with changes in membrane thickness. Mitochondria form large, dynamic networks that can exhibit distinct cellular distributions, including highly elongated (i.e., hyperfused) or fragmented networks. We previously demonstrated that these distinct network morphologies are associated with significantly distinct membrane ultrastructures that vary in inter- and intra-membrane spacing, curvature, and orientation^14^. We set out to understand if a similar connection exists between bulk mitochondrial morphology (i.e., elongated versus fragmented) and mitochondrial membrane thickness. We performed cryo-fluorescence microscopy to classify the network morphology of each cell prior to cryo-FIB milling and cryo-ET acquisition (***Supplemental Figure 3A)***^14^. We calculated the average membrane thickness across mitochondrial membrane surfaces and observed no statistically significant differences in the thickness of the OMM or IMM (and IMM subcompartments) based on network morphology (***Supplemental Figure 3B***), in contrast with previous work showing differences in inter-membrane spacing and curvature^14^.

Previous studies using *in vitro* reconstituted membranes demonstrated that membrane rigidity (i.e., the resistance to curvature) increases with thickness^19^. We observe significant variability in the curvature of the IMM, with the crista junction and the crista “tip” exhibiting the highest degree of curvature relative to the crista body and IBM (***Figure 4A***)^14^. Therefore, we wondered whether increases in membrane curvature are associated with thinner membrane regions in organellar membranes within the native cellular environment. To test this, we calculated the curvature of all triangles within the IMM surface mesh reconstructions and partitioned the curvature values into separate quantile ranges for comparison (0-0.5, 0.5-0.9, 0.9-0.95, 0.95-0.99, 0.99-1). Interestingly plotting the membrane thickness for each triangle within the curvature quartiles showed that triangles with the highest curvature (0.99-1 quantile) were associated with significantly thicker membranes relative to medium and low curvature regions (***Figure 4B***).

**Figure 4.**
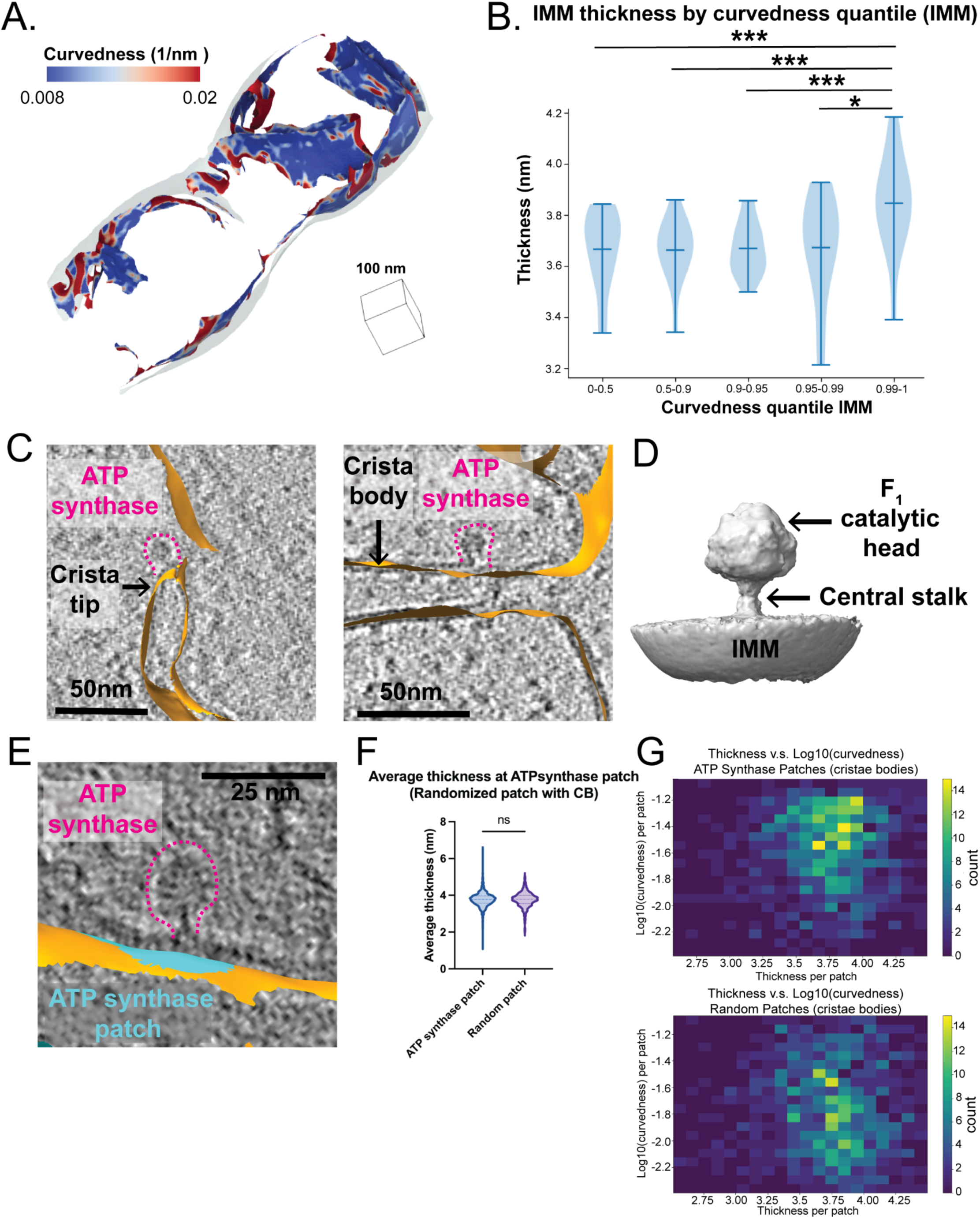
Membrane thickness is positively correlated with membrane curvature. A. Representative membrane surface reconstruction of mitochondria colored by IMM curvedness. B. Quantification of IMM thickness by curvedness quantile shows that higher curvature significantly correlates with thicker membrane. P values from Mann-Whitney U test are indicated. * P < 0.05, *** P < 0.005. C. ATP synthases (highlighted by pink dashed lines) are located at the IMM (orange) with distinct curvature. The left panel shows an ATP synthase positioned at a curved cristae tip, and the right panel shows one on a flat crista body. D. 3D subtomogram average structure of ATP synthase monomer positioned on IMM with its F1 catalytic head and central stalk highlighted by arrows. E. ATP synthase (outlined by pink dashed lines) and the corresponding membrane patch (cyan) representing its local ‘footprint’ on IMM (orange) F. Quantification of average thickness of ATP synthase patches and randomized patches on cristae bodies for each mitochondrion. ATP synthase patches show no significant difference in thickness compared to randomized patches. Quantification of ATP synthase patch number n = 984 and randomized patch number n = 984 is shown. G. 2D histogram of average thickness and log10 (curvedness) for ATP synthase patches and randomized patches on cristae bodies. ATP synthase patches show a positive correlation between thickness and curvature.

Given that ATP synthase dimers have the capacity to induce high degrees of membrane curvature^20^, we wondered whether the presence of ATP synthase in our datasets was associated with local changes in membrane thickness. In species like *Chlamydomonas reihardtii*, ATP synthase is primarily organized as dimer rows enriched at cristae ‘tip’ regions. In our dataset, we observe clear densities for ATP synthase that are assembled in a mixture of dimers and monomers, both in the cristae ‘tip’ and the cristae body regions (***Figure 4C***). To ask whether these ATP synthase complexes are associated with regions of high curvature and membrane thickness, we manually picked 8,993 ATP synthase monomers within our data and performed subtomogram averaging to further refine the position and orientation of these particles. This resulted in a 13 Å structure resembling ATP monomers from previous reports^21^ (***Figure 4D, Supplemental Figure 3C***). We applied a ‘patch-based’ analysis^22^ to subselect local patches of surface mesh reconstructions that correspond to the membrane ‘footprint’ of ATP synthase on the cristae body region of IMM, where ATP synthases are predominantly localized. In brief, this involved identifying the nearest IMM surface triangles of each ATP synthase particle coordinate and extracting the surrounding triangles within a radius of 120 Å of those nearest triangles (***Figure 4E***). Calculating membrane thickness locally at each ATP synthase patch region revealed no significant differences in membrane thickness compared to patches generated at random positions within the crista body (***Figure 4F***). However, in the context of curvature, we observed that those ATP synthase patches at high curvature regions were also associated with thicker membranes, compared to what would be expected from random chance (***Figure 4G***). This suggests that individual ATP synthase particles associated with high-curvature regions are also found in regions of greater membrane thickness.

### Local changes in membrane thickness are co-localized with membrane-associated proteins that exhibit unique density line scan profiles

Unique to our approach is the ability to accurately identify local variability in membrane thickness on a locally-averaged, per-triangle basis (***Figures 2A* *& 5A***). Within our surfaces, we isolated the local patches that exhibited the most increased or reduced membrane thickness relative to the surrounding areas (***Figure 5A*).** Mapping these local regions back to their locations within the tomogram revealed that they often co-localize with large, membrane-embedded complexes (***Figure 5B***). Surprisingly given our previous analysis, we observe ATP synthase localizing to thicker regions (***Figure 5B***). Strikingly, we detect several macromolecules localized to regions of thinner membrane (***Figure 5B***). Although it is challenging to unambiguously identify all these macromolecules based on their size and location within the crista, a subset likely represents components of the oxidative phosphorylation machinery. To further investigate the unique structural properties of these macromolecules, we extended our patch-based^22^ density scans to encompass regions beyond the IMM, extending into the mitochondrial matrix. All line scans show the characteristic double Gaussian peaks of the IMM, with additional peak profiles observed at greater distances away from the IMM surface (***Figure 5C***). For ATP synthase, this corresponds to an additional broad peak spanning from 7 nm to 17 nm, corresponding to the diameter of the F1 catalytic domain. Comparing the average of several line scans of ATP synthase with those generated using randomized patches along the IMM caused this peak to disappear (***Figure 5D***). This suggests that this patch-based line scan approach can uniquely identify distinct signatures of macromolecules, paving the way for future applications aimed at subclassifying distinct membrane-associated molecules in cells.

**Figure 5.**
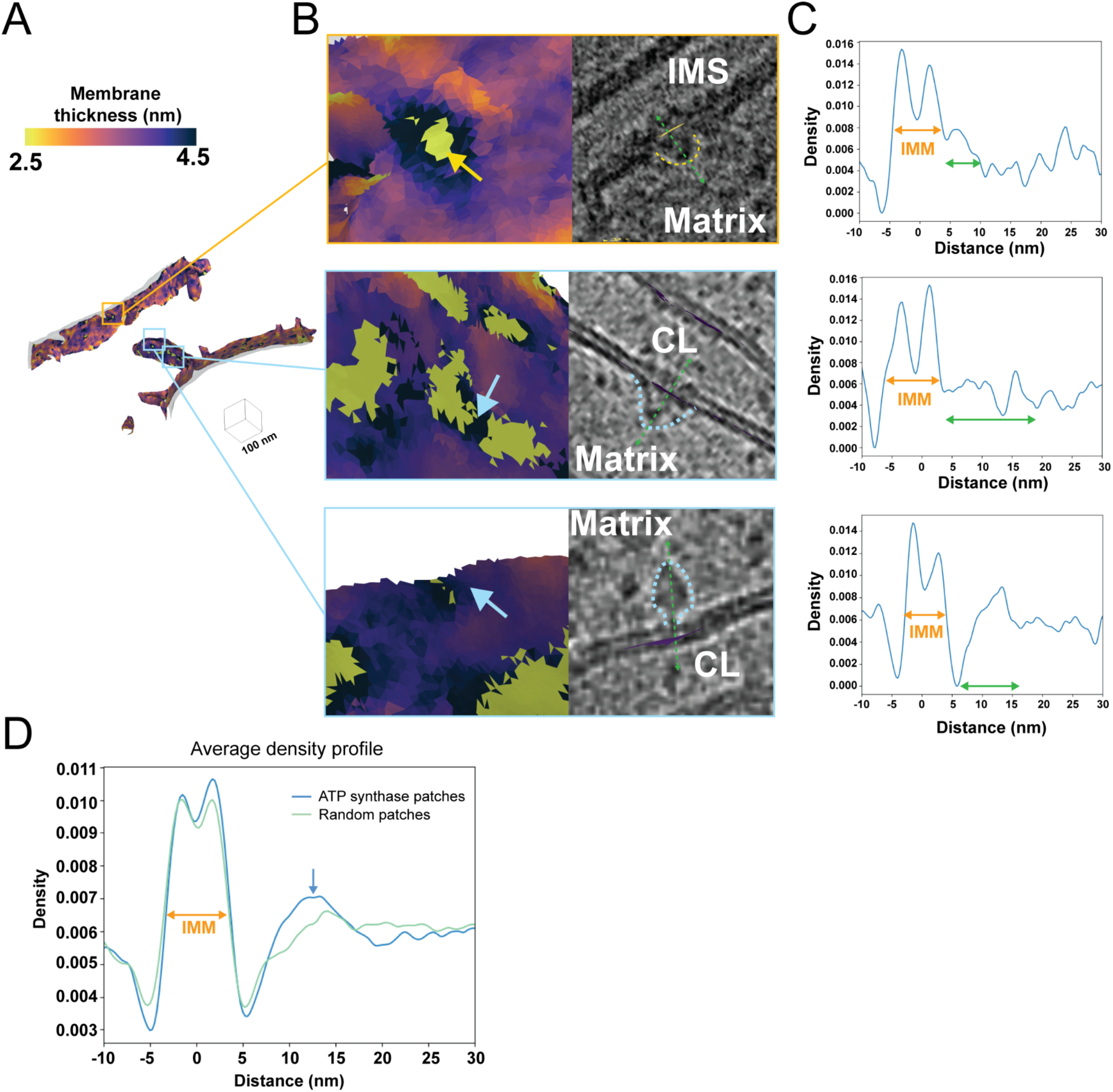
Local variations in membrane thickness correspond to the presence of membrane-associated proteins. A. Representative membrane surface reconstruction of mitochondria colored by IMM membrane thickness. B. Enlarged images of the corresponding boxed regions shown in (A) reveal the local membrane thickness and the co-localized membrane-associated macromolecules (outlined by dashed lines). C. Line scan profiles along the green dashed arrow line in (B) of membrane-associated macromolecule patches show double Gaussian peaks corresponding to IMM (orange arrows), along with additional peaks representing protein macromolecules (green arrows). D. The averaged line scan density profile across all ATP synthase patches (blue) shows a distinct peak compared to randomized patches (green).

## Discussion

The diversity of organelle membranes has been extensively studied through lipidomics^23,24^, proteomics^25,26^, light^27^ and electron microscopy^28,29^. However, recent advances in cryo-electron tomography have enabled the direct visualization of these membranes within cells with higher resolution than ever before. We describe an automated and robust method to measure organelle membrane thickness directly from observed density in cellular cryo-electron tomograms. A substantial body of work has used contour-based density scanning in single-tilt images of *in vitro* membrane bilayers^9^; our method advances these measurements into three dimensions within cells by taking advantage of our previously developed tools to automatically model membranes as triangle meshes in tomograms^14^. These surface meshes serve as a scaffold to perform hundreds of thousands of voxel-based density line scans across entire organelles.

A key advantage of this method is that it is agnostic to the approach used for generating the initial voxel-based segmentation, due to the combination of the surface mesh model and the direct sampling of the original tomogram for thickness measurements. Various techniques exist for isolating or segmenting membranes from cryo-tomograms into binarized volumes that can subsequently be attributed to specific organelles. These range from watershed transforms^30^, to computer vision-based methods such as tensor voting (e.g., TomoSegMemTV^31^, ColabSeg^32^), as well as more recent approaches using 2D/3D U-net architectures (e.g., EMAN2^33^, Membrain-seg^15^, Deepfinder^34^). To address segmentation challenges posed by low signal-to-noise ratios, many of these methods prioritize voxel intensity connectivity to generate visually complete membrane segmentations. While appropriate for visualization, this can lead to variations in membrane width. The expanded Surface Morphometrics pipeline overcomes these limitations by directly performing voxel-based line scans on the cryo-tomography. Additionally, tomograms can be interchanged with different binning or additional post processing with denoising or contrast enhancement algorithms as long as tomogram relative dimensions remain consistent– allowing for rapid adaptation of workflows.

We applied our method to analyze organelle membranes within mouse embryonic fibroblasts and identified statistically significant differences across various physiological membranes. We capture significant differences between the membranes of mitochondria in which the OMM has a thickness of 3.2±0.10 nm, thinner than the IMM, which has a thickness of 3.6±0.07 nm (***Figure 2***). The inner mitochondrial membrane is one of the most protein-rich membranes in the cell, having been reported as containing 60-70% protein content by mass^35^. In contrast, the outer mitochondrial membrane is reported to have a much lower protein content – 45% protein content by mass^36^. In addition to protein differences, the OMM and IMM also have distinct lipid compositions with the IMM being enriched in cardiolipin promoting higher curvature and stabilizing large protein complexes^37^. Beyond these differences, the observed vesicles (3.6 nm) were significantly thicker than both the ER and the OMM. We did not distinguish different vesicle types (which would typically need specialized CLEM approaches), and thus we can’t explain why vesicles have thicker membranes. It may be due to specific membrane-associated cargo or due to specific lipid content. Further studies of vesicles with specified origins, as well as with vesicle-originating organelles such as the plasma membrane and the Golgi apparatus, will help to differentiate these possibilities, as will studies of *in vitro* vesicles with defined lipid and protein composition.

Beyond differences in thickness between organelles, we detected local variation in thickness within physiologically distinct organelle subcompartments. In the inner mitochondrial membrane, we identified variation between the inner boundary membrane, crista junctions, and crista bodies. These have sequentially increasing thicknesses, with the crista body significantly thicker than either of the other two compartments (***Figure 3***). Given the continuous membrane bilayer connecting these subcompartments, this local variation is likely due to a combination of local lipid enrichment and differences in protein content within the different subcompartments. In contrast, when comparing the differences between rough and smooth ER, the small apparent thickness difference was not statistically significant, though this may be due to the limited number of surfaces measured (smooth ER: n = 5 surfaces, rough ER: n= 4 surfaces). ER structures are diverse^38,39^ and we anticipate that differences in thickness may often be found in physiologically distinct membrane subcompartments with local enrichments of both lipid and protein factors.

We also found enhanced bilayer thickness in the most curved segments of the IMM (***Figure 4***). This is in sharp contrast to *in vitro* biophysical studies, which reported that membrane resistance to curvature increases quadratically with bilayer thickness - the “thicker sandwich” is harder to bend^19^. This increase in thickness at the top 1% most curved triangles of the highest degrees of curvature is biophysically unfavorable, as thicker membranes are more rigid. This suggests that additional forces, such as membrane-shaping proteins, may contribute to the increase in membrane thickness locally at these high-curvature regions.

We reason that this difference from biophysical principles must be due to either specific lipids (cardiolipin, in particular, may increase both curvature and thickness together)^40^ or membrane-bending proteins involved in generating that increased thickness. To disentangle these possibilities, we attempted to evaluate the role of ATP synthase, one of the best characterized proteins involved in driving membrane curvature in cristae. Consistent with its role, we show that regions of extremely high curvature in the IMM are associated with ATP synthase molecules. However, when comparing membrane regions in the presence and absence of ATP synthase we did not identify any significant changes in membrane thickness. One potential explanation is that within our dataset we observe a mixture of ATP synthase monomers and dimers, and analyzing this heterogeneous population may obscure differences in membrane thickness.

Although ATP synthase did not significantly colocalize with membrane thickness, we discovered that many of both the thickest and thinnest sections of membranes are associated with large membrane-associated protein complexes (***Figure 5***). This aligns with our current understanding of protein-lipid interactions, which proposes that specific microdomains help stabilize and regulate protein complexes, such as the respiratory complexes^41^. This is suggestive that membrane bilayer variation may be a sensitive signature to help identify challenging-to-find membrane proteins within tomograms. In support of this concept, we extended our line scan approach beyond the lipid bilayer into the space around the membrane and showed that complexes like ATP synthase exhibit distinct line scan profiles. This opens the door to incorporating membrane mesh-guided density sampling as an assistive tool in protein localization and identification within tomograms.

In summary, our model-guided density-based approach to measuring membrane thickness reveals variation between organelles and within distinct organelle subcompartments, as well as within organelles in correlation with ultrastructural features such as curvature. Furthermore, this new and robust tool for measuring bilayer thickness has been incorporated into the Surface Morphometrics pipeline, enabling measurement of thickness from segmented tomograms in a fully automated manner in concert with other features such as membrane-membrane distances, curvature, and orientation. In this way, it will be straightforward for microscopists to measure the association of bilayer thickness with features such as membrane contact sites or curvature-separated subcompartments. We have already demonstrated the value of an early version of this approach in recent collaborative work studying ER-budded replication organelles in arbovirus-infected cells^42^. We look forward to seeing the discoveries made by other researchers using this new tool in the Surface Morphometrics pipeline.

## Acknowledgments

We thank Bill Anderson and William Lessin at The Scripps Research Institute Hazen cryo-electron microscopy facility for microscope support. We thank Jean-Christophe Ducom and Lisa Dong at The Scripps Research Institute for computational support. We also thank R. Luke Wiseman, David DeRosier, Reika Watanabe, Kelsey C. Martin, and all other members of the Grotjahn Lab for their critical input on the manuscript. M.M is supported by the ARCS (Achievement rewards for college scientists) foundation. B.A.B. is supported by the Collins Medical Trust. D.A.G. is supported by The Pew Scholars Program, Nadia’s Gift Foundation Innovator Award of the Damon Runyon Cancer Foundation (DRR-65-21) and the National Institutes of Health (NIH) grant RF1NS125674. This work used equipment supported by NIH grant S10OD032467.

## Author Contributions

**M. Medina:** Conceptualization, Data curation, Formal analysis, Investigation, Methodology, Validation, Visualization, Writing - original draft, Writing - reviewing and editing.

**Y.-T. Chang:** Conceptualization, Formal analysis, Investigation, Methodology, Software, Validation, Visualization, Writing - original draft, Writing - reviewing and editing.

**H. Rahmani:** Data curation, Investigation, Methodology, Validation, Writing - reviewing and editing.

**D. Fuentes:** Data curation, Investigation, Methodology.

**B. A. Barad:** Conceptualization, Formal analysis, Investigation, Methodology, Project Administration Methodology, Software, Supervision, Validation, Visualization, Writing - original draft, Writing - reviewing and editing.

**D. A. Grotjahn:** Conceptualization, Funding acquisition, Methodology, Project administration, Resources, Supervision, Validation, Visualization, Writing - original draft, Writing - reviewing and editing.

## Methods

### Preparation of vitrified mouse embryonic fibroblasts on cryo-EM Grids

Mouse embryonic fibroblasts expressing mitochondrially-localized GFP (MEF^mtGFP^)^43^ were cultured in Dulbecco’s Modified Eagle Medium + GlutaMAX (Gibco) additionally supplemented with HiFBS (10%) and glutamine (4 mM) on fibronectin-treated (500 ug/ml, Corning) and UV sterilized R ¼ Carbon 200-mesh gold electron microscopy (EM) grids (Quantifoil Micro Tools). After 15-18 hours of culture, MEF^mtGFP^ cells were plunge-frozen in a liquid ethane/propane mixture using a Vitrobot Mark 4 (Thermo Fisher Scientific). The Vitrobot was set to 37° C and 100% relative humidity and blotting was performed manually from the back side of grids using Whatman #1 filter paper strips through the Vitrobot humidity/temperature chamber side port. The Vitrobot settings used to disable automated blotting apparatus were as follows: Blot total: 0, 2; Blot force: 0, 3; Blot time: 0 seconds.

### Cryo-fluorescence microscopy and Mitochondria Network Morphology Scoring

Fluorescence and bright-field tiled image maps (atlases) of EM grids containing vitrified cellular samples were acquired with a Leica CryoCLEM microscope (Leica) were collected using Leica LAS X software (25 um Z stacks with system optimized steps, GFP channel ex: 470, em: 525). Z stacks were stitched together in LAS X navigator to provide a single mosaic of maximum projections of the GFP signal, enabling rapid identification of the bulk mitochondrial morphology for each cell. For classification of mitochondrial network morphologies, max projections of individual tiles within fluorescence atlases of MEF^mtGFP^ cells were randomized and blinded. Five researchers classified the cells as containing primarily elongated or fragmented mitochondria. Atlases were then exported from LAS X and loaded into MAPS software (Thermo Fisher Scientific) for fluorescence guided milling.

### Fluorescence Guided Milling

Cryo-Focused Ion Beam (FIB) milling of lamellae was performed using an Aquilos dual-beam cryo-FIB/SEM instrument (Thermo Fisher Scientific) operated by xT software (Thermo Fisher Scientific). The fluorescence atlases were overlaid and aligned to an SEM atlases of the same grid to target milling of MEF^mtGFP^ cells with distinct mitochondrial network morphologies, as determined during blind classification (described above). MEF^mtGFP^ cell targets were chosen based on their position within grid squares, the thickness of the ice in their vicinity, and their bulk morphology as assessed by the GFP fluorescence channel. Prior to milling, EM grids were first coated with an organometallic platinum layer using a gas injection system (GIS) for 3-4 seconds using an automation script ^44^, followed by a layer of platinum sputter. Targeted cells were milled using an automated cryo preparation workflow^45^ both methods used xT software with MAPS (Thermo Fisher). Minimal SEM imaging for monitoring was done at 2keV to ensure thin lamella generation while avoiding radiation damage. Upon completion of final polishing of lamellae an 8 sec layer of platinum sputter was added to deposit platinum (bead) fiducials for downstream tilt series alignment. A total of 2 grids were milled for further tomography analysis.

### Tilt Series Data Collection

EM grids containing lamellae were transferred into a 300keV Titan Krios microscope (Thermo Fisher Scientific), equipped with a K3 Summit direct electron detector camera (Gatan), and a BioQuantum energy filter (Gatan). Individual lamellae were montaged with low dose (1 e^-^/Å^2^) at high magnification to localize cellular regions containing mitochondria, which were identified by their distinctive inner and outer mitochondrial membranes. Data was collected to maximize the number of non-overlapping fields of view containing mitochondria, with no targeting of specific observed membrane ultrastructure. Tilt series were acquired using parallel cryo-electron tomography (PACE-tomo) (Eisenstein, Yanigasawa et al. 2023), which is a set of Python-based SerialEM (Mastronarde, 2005) allowing multiple tilt series collection in parallel on the same lamella using beam shift. Tilt series were acquired at magnification 53,000x with a pixel size of 1.662 Å and a nominal defocus range between (−4 to −8 µm). Data collection was done in a dose symmetric scheme with 2° steps between -60° and +60° centered on −11° pretilt. Data was collected with dose fractionation, with 10 0.3001 e/Å^2^ frames collected per second. The total dose per tilt was 3.0 e/Å^2^, and the total accumulated dose for the tilt series was under 123 e/Å^2^.

### Tilt series processing and reconstruction

Dose fractionated tilt series micrograph movies underwent CTF estimation and motion correction in Warp^46^ and combined into averaged tilt series for alignment. Fractionated tilt series then were aligned using bead alignment using the post polish platinum fiducials in etomo ^47^. In some cases, the coverage of the platinum fiducials on the tilt series position was not amenable for bead tracking and patch tracking in etomo was used with 4 times binning and 36 binned pixel patches. Resulting contours were manually curated to remove poorly aligning patches, and the remaining contours were used for alignment and reconstruction with Warp^46^ back projection into 4, 6, and 8 times binned tomograms. Fiducial platinum “beads” were erased using a BoxNet model in Warp. Tomogram thicknesses ranged from 90 nm to 220 nm.

### Membrane tracing, voxel segmentation, and surface generation

All reconstructed tomograms with eight times binning (voxel dimensions:13.30Å x 13.30Å x 13.30Å) were processed by Membrain-Seg^15^, which is an advanced machine learning software based on U-Net architecture for tracing and segmenting cellular membranes. The binarized volumes of the traced membranes were then input into AMIRA (Thermo Fisher Scientific) for manual curation. The mitochondrial membranes (OMM and IMM) and ER membranes were designated as different labels using the 3D magic wand tool. Manual clean up of organelle membranes was performed using the 2D paintbrush tool. Final voxel segmentations were confirmed by visual inspection in AMIRA with comparison to original tomogram. The voxel segmentation membrane label files were then exported from AMIRA and input into Surface Morphometrics^14^. The labels of each membrane voxel segmentation were reconstructed as smooth surface meshes using the “segmentation_to_mesh.py”. The surfaces were generated with a maximum of 200,000 triangles, a reconstruction depth of 8, and an extrapolation distance of 1.3 nm. Curvature estimations of triangulated surface meshes were run using “run_pycurv.py. Inter and intra-organelle distances were measured using “membrane_distance_orientation.py”.

### Thickness measurement

For each triangle in a surface mesh, the density in the cryoEM map was interpolated at along the normal vector at 0.25 nm increments ranging from 10 nm below the triangle to 10 nm above to produce a “line scan” revealing the electron density normal to the surface, revealing the areas of increased density corresponding to the head groups of each leaflet of the phospholipid bilayer. These linescans are generated in the ‘thickness_scan.py’ script. These individual scans are very noisy due to the low signal-to-noise inherent to tomography data; in order to generate scans that could consistently fit, we applied two averaging strategies. For global thickness measurement, all line scans in a surface were averaged before fitting with a dual gaussian distribution. For local measurements, each triangle’s line scan was averaged with all neighboring triangles within 12 nm of the original triangle, with weighting that decreased with the distance from the central triangle by the following formula: 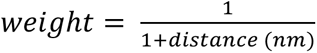. In this way, closer triangles counted more for the assessment of thickness, minimizing the loss of local detail while enhancing signal to noise. With these averaged line scans, the phospholipid head groups were modeled by fitting a dual gaussian distribution to the line scan. The measured thickness was determined by the difference in the means of the two gaussians. These thickness calculations were accomplished with the ‘thickness_plots.py’ script. For each surface, the median thickness was calculated using weights corresponding to the area of each triangle, accounting for triangle size variation. The mean and standard error-based 95% confidence interval of these per-surface measurements were reported and all statistical comparisons of different surface types used the Mann-Whitney U-test^48^. Statistics were generated using the ‘statistics_stats.py’ script.

### Patch based analysis

To analyze the local environment of ATP synthase, we defined ATP synthase-associated patches as regions on the IMM surface where ATP synthase particles localize. ATP synthase coordinates were obtained from the starfiles corresponding to each tomogram. The IMM surface coordinates and thickness for each mitochondrion were from triangle graph files (.gt) generated by Surface Morphometrics pipeline. To identify patches for each mitochondrion, we first located the nearest IMM surface triangle to each ATP synthase particle using a k-dimensional tree Python function. To avoid cross-assignment ATP synthase from other mitochondria in the same tomogram, we excluded any nearest IMM triangles located farther than the height of an ATP synthase particle (24 nm). The remaining nearest IMM surface triangles were designated as “patch centers.” Around each patch center, we searched for triangles within a 12 nm radius to define the ATP synthase-associated patch. Randomized ATP synthase-associated patches for each mitochondrion were generated based on the following criteria: (1) the number of randomized patches matched the number of ATP synthase-associated patches, and (2) the distances between randomized patch centers were greater than 12 nm. This process was performed using “find_IMM_patches_for_ATP_synthase.py”. The average thickness for each ATP synthase-associated patch and each randomized patch were calculated by “average_thickness_calculation_per_patch.py” and visualized as a violin plot. The Mann–Whitney U test was applied to assess the statistical significance of differences. The generation of the violin plot and statistical test were performed using Prism. The average curvature for each ATP synthase-associated patch and each randomized patch was calculated by “average_curvature_calculation_per_patch.py”. The average thickness and log_10_ curvedness are plotted as a 2-dimensional histogram by “2dhist_curvedness_thickness.py”.

In order to obtain the average ATP synthase line scanning density profile, we extracted ATP synthase-associated patches from multiple mitochondria as individual patch surfaces, preserving the coordinates and normal vectors of each surface triangle, using the script “extract_single_patch.py”. Before performing line scanning, the normal vectors of the surface triangles in each patch were curated to ensure they pointed in the same direction as the vector from the patch center to the corresponding ATP synthase particle center. These curated patch surfaces were then correlated with the tomogram and served as the reference for the line scanning process. For each ATP synthase, a line scanning density profile was generated by sampling tomogram intensity values along the direction of the curated normal vectors. The scan extended from −10 nm (toward the inner membrane space) to +30 nm (toward the matrix) with 0.25 nm steps. The average ATP synthase line scanning density profile was then obtained by averaging the intensity profiles across all ATP synthases. To obtain the line scanning profile for the other membrane-associated proteins, we picked the protein candidates as particles in ArtiaX and identified the corresponding patches with 12 nm radius by the same approaches as ATP synthase. We performed line scans ranging from −10 to 30 nm on individual membrane-associated protein patches, using particle-curated vectors to obtain the density profile for each membrane-associated protein.

### Subcompartment analysis

Rough ER and smooth ER were separated on entire surfaces by visual inspection of the tomogram, identifying ribosome-bound ER membranes. For sub compartment analysis of IMMs, the inner membrane surface was subdivided based on distance from the OMM relative to the mode distance measured for each surface, which corresponds to average spacing between the OMM and the IBM. All triangles less than 4nm beyond this distance were classified as IBM. Triangles between 4nm and 14nm beyond this distance were classified as junctions. All triangles more than 4 nm beyond this distance were classified as crista bodies.

### Subtomogram averaging of mitochondrial ATP synthase complexes

ATP Synthase complexes were picked using a two-point directional picking strategy on a larger dataset of 43 tomograms from MEF^mtGFP^ cells that were either treated with Tg (500nm), ionomycin (1uM) or Vehicle. Tomograms of binning 4 (6.65 A) were post-processed with a median filter in z direction (one iteration and kernel size of 11 pixels) and an mtf filter and used to manually particle pick prohibitin complexes using the software package i3^49^. These particle picks were then converted to star files, and then warp2dynamo package was used to create dynamo tables^50^. These particles were refined in dynamo for 5 cycles using the global refinement preset, and then the rotation angles were randomized using *dynamo_table_randomize_azimut*h to avoid aligning all of the particles on the missing-wedge artifact. This randomized table was then converted to star files using warp2dynamo package and particles were extracted using Warp and refined in Relion (relion_autorefine) to achieve the resolution of 19 Å^51^. Half-maps were post-processed and refined in M and reached the resolution of 12.9 Å^52^. ArtiaX module in Chimerax was used to visualize these particles in the original tomograms^53,54^.

### Data and Code availability

All tilt series, reconstructed tomograms, voxel segmentations, and reconstructed mesh surfaces used for quantifications were deposited in the Electron Microscopy Public Image Archive (EMPIAR) under accession codes EMPIAR-XXXXX. All subtomogram averages were deposited in the Electron Microscopy Data Bank (EMDB) under accession codes EMDB-XXXX. All scripts used for Surface Morphometrics are available at https://github.com/grotjahnlab/surface_morphometrics).

