## Supplemental Figures for "Surface morphometrics reveals local membrane thickness variation in organellar subcompartments"

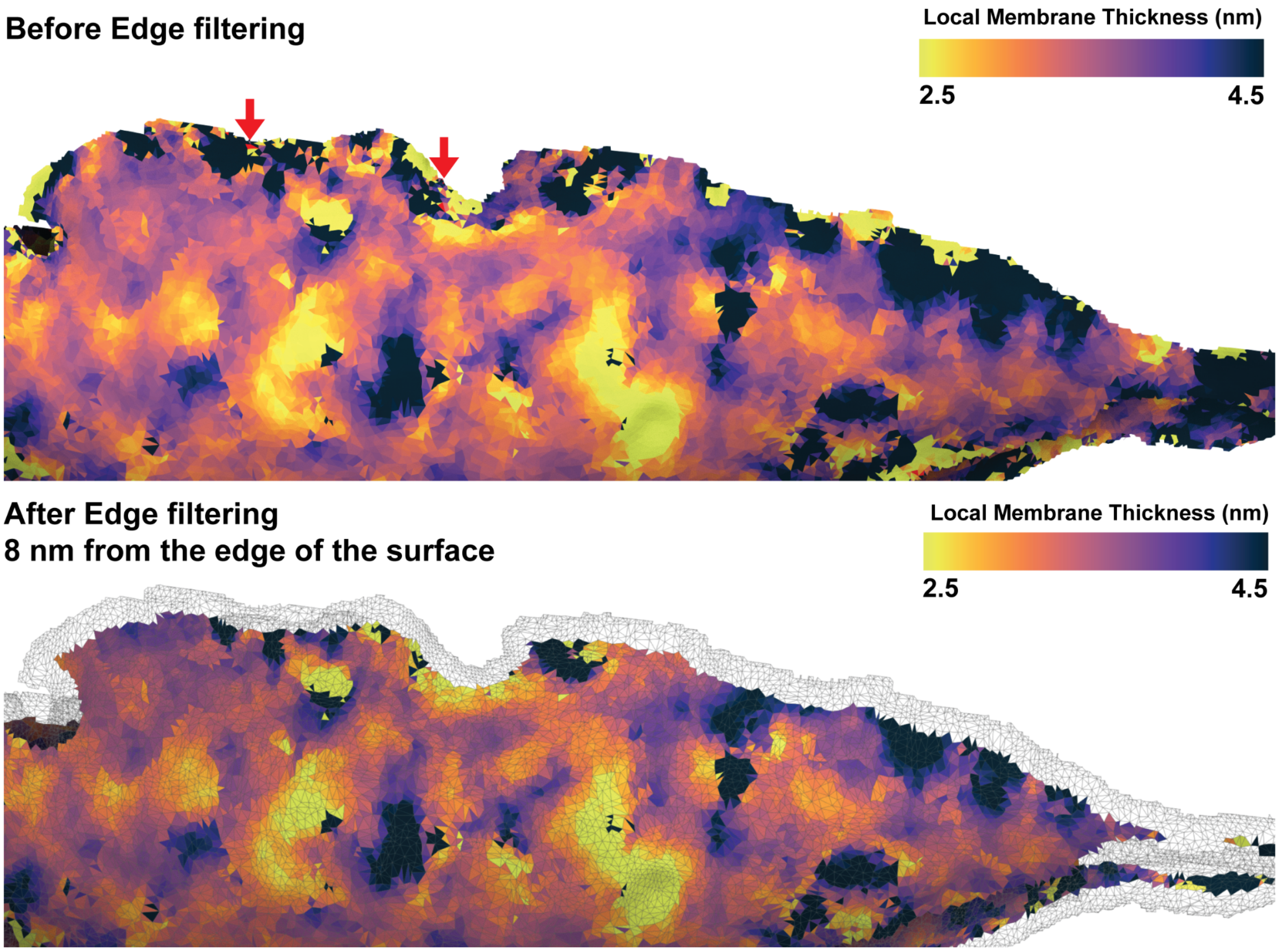


**Supplemental Figure 1. Edge filtering to improve measurement robustness.**

To remove artifactual measurements at the edge of surfaces, the 8 nm nearest to the edge of the membrane mesh are artificially removed. Arrows highlight areas where measurements could not be made at all.

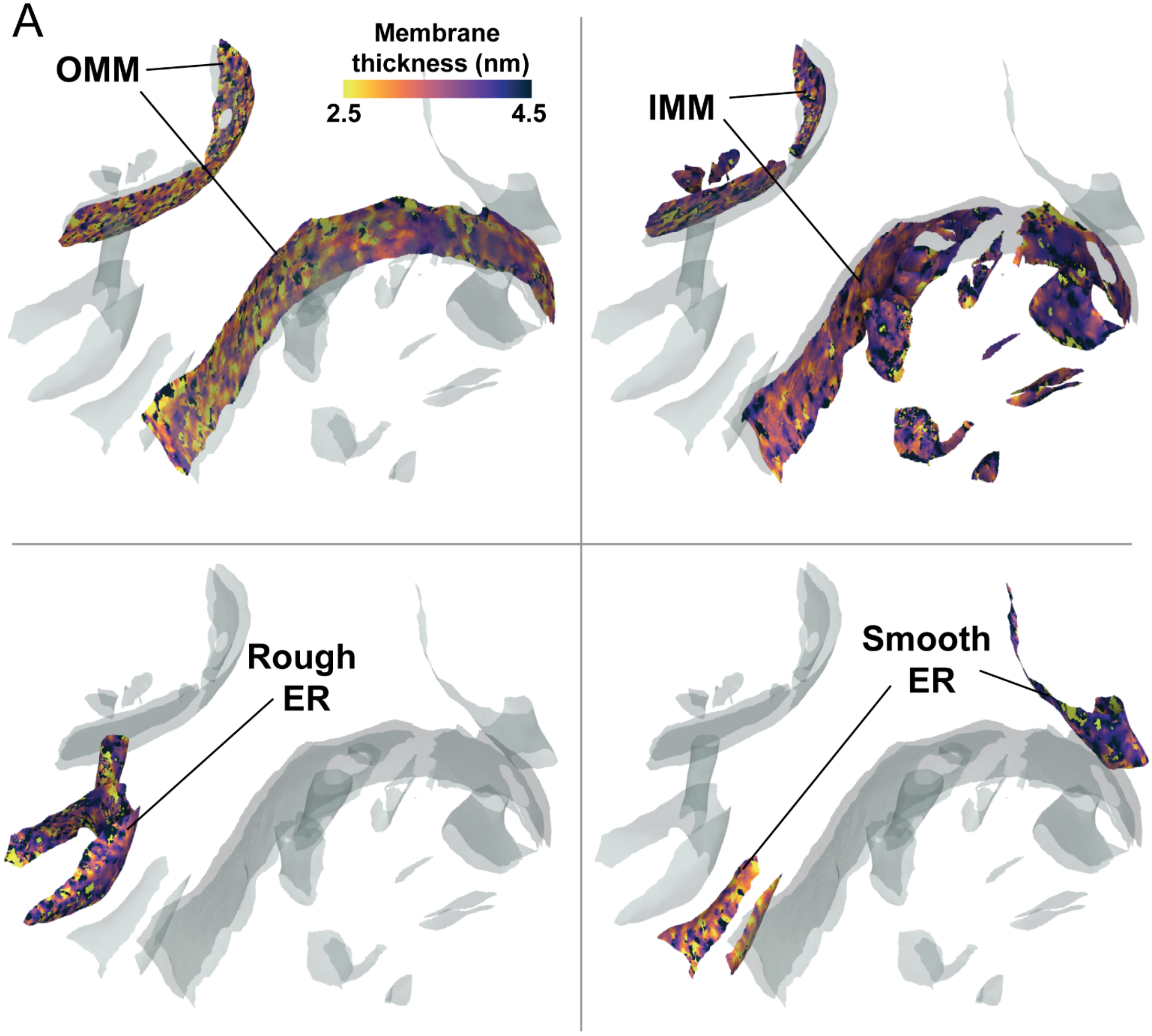
A
Membrane
thickness (nm)
2.5
4.5
OMM
IMM
Rough
ER
Smooth
ER

**Supplemental Figure 2. Gallery of local thickness variations in organelles**

Local thickness variations of individual physiological organelles are observed within a single tomogram, highlighting the variations both within and between organelles.

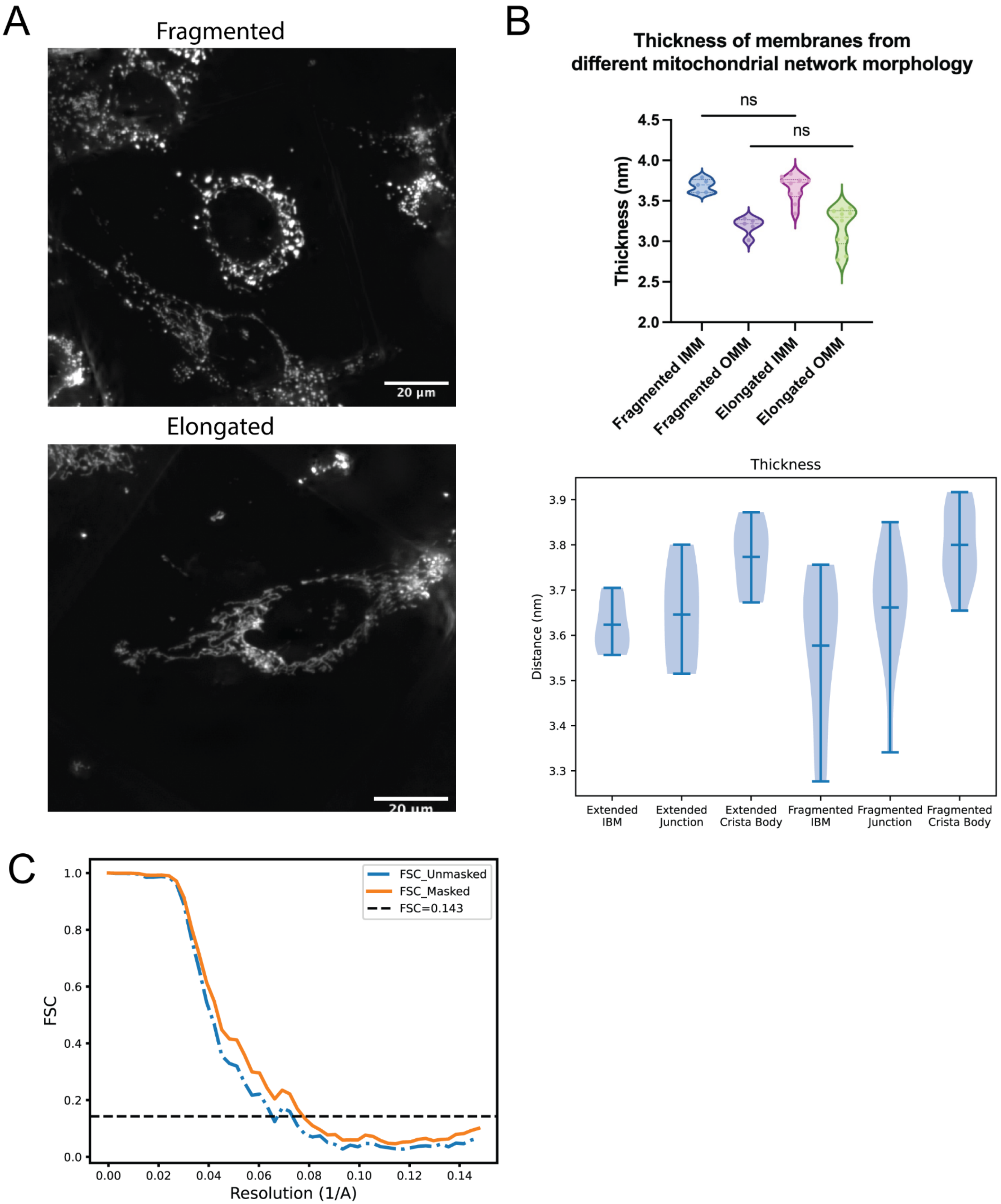
A
Fragmented
20 µm
Elongated
20 µm
B
Thickness of membranes from
different mitochondrial network morphology
ns
ns
Thickness (nm)
4.5
4.0
3.5
3.0
2.5
2.0
Fragmented IMM
Fragmented OMM
Elongated IMM
Elongated OMM
Thickness
Distance (nm)
3.9
3.8
3.7
3.6
3.5
3.4
3.3
Extended IBM
Extended Junction
Extended Crista Body
Fragmented IBM
Fragmented Junction
Fragmented Crista Body
C
FSC_Unmasked
FSC_Masked
FSC=0.143
FSC
1.0
0.8
0.6
0.4
0.2
0.0
0.00
0.02
0.04
0.06
0.08
0.10
0.12
0.14
Resolution (1/A)

**Supplemental Figure 3. Membrane thickness is not correlated with network morphology.**

A. Cryo-fluorescence microscopy of elongated and fragmented mitochondrial networks in $MEF^{mitoGFP}$ cells.

B. Violin plots displaying membrane thickness across different mitochondrial membranes and IMM subcompartments based on cellular mitochondrial network morphologies.

C. FSC curve of ATP synthase structure.

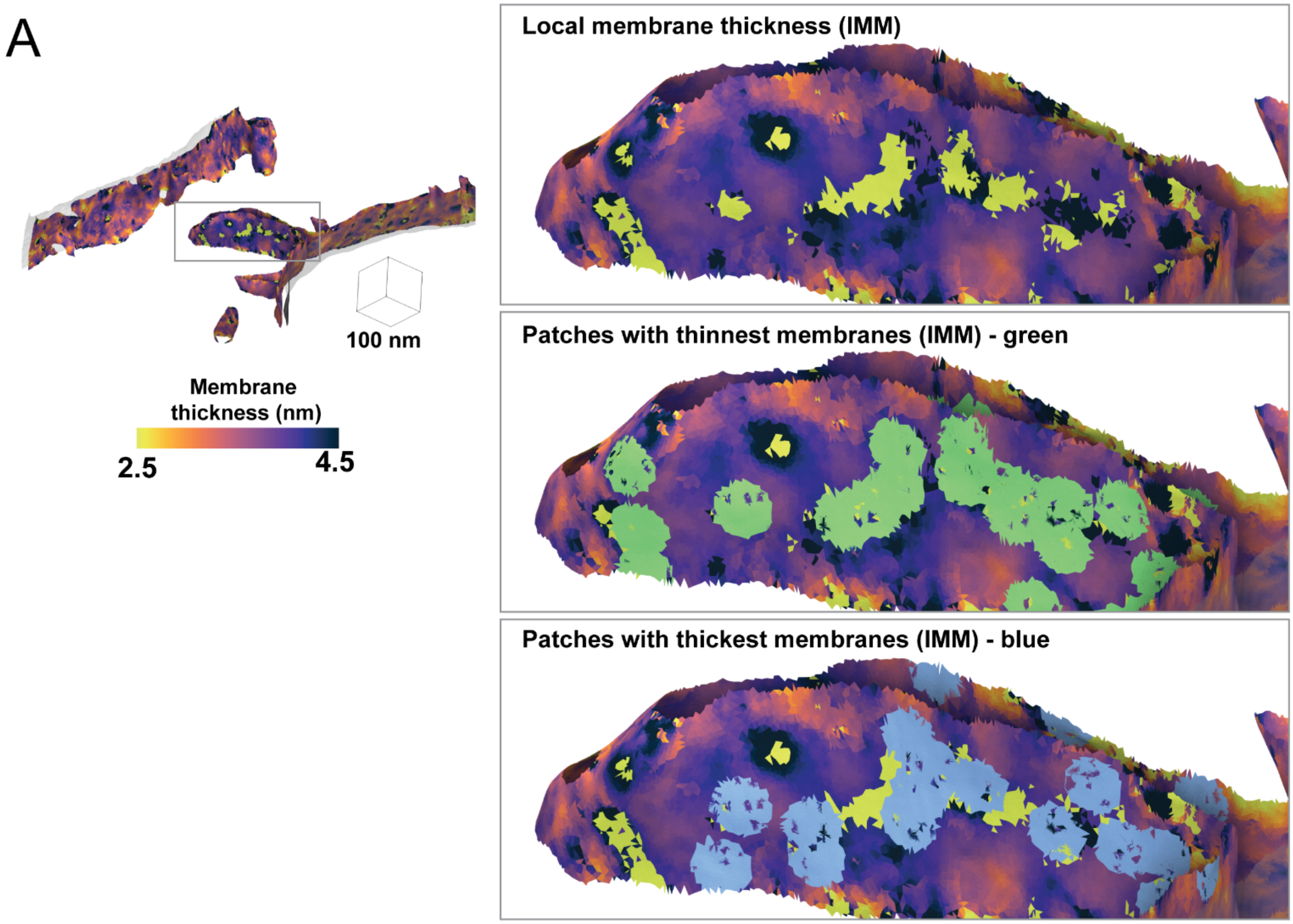


**Supplemental Figure 4. Extreme thickness measurements reside in the mitochondria cristae.**

The per-triangle local thickness variation within a tomogram can be used to generate patches with the 50 thinnest (green patches) and thickest (blue patches) triangles within each tomogram; these patches can be used for visually assessing protein content at these locations.
